# Deep phenotypic profiling uncovers cryptic effects of antifilarial drugs

**DOI:** 10.64898/2026.02.12.705610

**Authors:** Kaetlyn T. Ryan, Nathalie Dinguirard, Clair R. Henthorn, Nicolas J. Wheeler, Mostafa Zamanian

## Abstract

The anthelmintics ivermectin, albendazole, and diethylcarbamazine are the backbone of mass drug administration (MDA) campaigns targeting human filariasis, yet their direct effects on parasites are still not fully defined or understood. The clinical effects of these drugs are stage dependent, resulting in effective clearance of circulating microfilariae but only limited activity against adult worms, a pattern that complicates disease surveillance and elimination efforts. Although molecular targets have been identified or proposed for some antifilarial drugs, their precise modes of action remain opaque, and conventional *in vitro* assays of motility or viability have generally failed to reflect pharmacologically relevant effects. There is growing evidence that cryptic phenotypes involving altered host-parasite interactions, including changes in parasite secretions, may help reconcile these discrepancies. Focusing on the causative species of lymphatic filariasis, we used high content imaging and quantitative mass spectrometry to enable deeper phenotypic profiling of drug responses in microfilariae and adult worms exposed to antifilarial compounds. In microfilariae, altered environmental conditions (temperature and salinity) lead to modest ivermectin effects on motility at therapeutic concentrations. In adult parasites, we show that drug responses vary with worm age and that different anthelmintics induce distinct changes in the secretory proteome. This improved phenotypic resolution advances our understanding of drug action in intra-host stages and highlights how antifilarial drugs can alter secretory cargo relevant to the detection of adult parasites that persist after drug treatment.

## Introduction

Lymphatic filariasis (LF) is a neglected tropical disease that is transmitted by mosquitoes infected with the parasitic nematodes *Wuchereria bancrofti*, *Brugia malayi*, and *Brugia timori* and causes severe chronic disability [1,2]. According to the World Health Organization (WHO), an estimated 51 million people are currently affected by LF, with 657 million at risk across 39 countries [3]. Current control and elimination strategies rely primarily on mass drug administration (MDA) using combinations of ivermectin (IVM), albendazole (ABZ), and diethylcarbamazine (DEC) to clear blood-circulating microfilariae (mf) and break the cycle of transmission [4,5]. While MDA programs have achieved significant progress toward reducing LF transmission and disease burden, several challenges remain. Antifilarial drugs require repeated annual treatment [6] and have limited efficacy against adult worms. These drugs can also cause adverse effects in regions co-endemic with other filarial diseases [7], and there is growing concern about the emergence of drug resistance, which has already been documented in closely related veterinary nematodes [8,9]. Finally, there is a critical need to improve methods to specifically detect the presence of surviving adult parasites in post-treatment surveillance [4,10].

A major factor contributing to these challenges is our incomplete understanding of how existing antifilarial drugs exert their effects. Although these compounds have been used for decades, their precise mechanisms of action and the full spectrum of their antiparasitic activity remain poorly defined. While broad classes of molecular targets have been identified for antifilarial drugs [11,12], how engagement of these targets translates into organismal phenotypes or stage-specific parasite clearance is not fully understood [13–15]. For example, ivermectin acts on glutamate-gated chloride channels (GluCls) but produces no overt *in vitro* phenotypes in mf at therapeutically relevant concentrations. Several studies have described *in vitro* inhibition of mf motility in response to IVM, but only at concentrations much higher than those required to clear parasites in the host [16–18]. This apparent disconnect between *in vivo* efficacy and *in vitro* effect has been partly reconciled by work showing that ivermectin inhibits mf secretory function through inhibition of protein and vesicle release [19–22]. Similar host-dependent or indirect mechanisms may underlie the actions of albendazole and diethylcarbamazine [23–25].

A clearer picture of these mechanisms could not only guide the discovery and development of more effective therapeutics but also improve how current drugs are deployed and monitored in elimination programs [26,27]. Drug responses are classically assessed through *in vitro* measures of parasite motility and viability [17,28–30]. Expanding the range of phenotypes measured across different environmental conditions, including those that reflect varying host states and directly or indirectly alter parasite secretory activity, would enable a more comprehensive assessment of drug action. Refining our understanding of these effects is also critical for identifying reliable molecular markers for surveillance applications in the context of treatment [31–34].

In this study, we used an image based phenotyping platform to assess *in vitro* drug responses in intramammalian life stages, both to validate established effects and to map broader response patterns relevant to the stage specificity of drug action. We examined how parasite culture conditions and age shape these responses, revealing that physiological context can sensitize parasites to antifilarial drugs at concentrations more aligned with therapeutic exposure. Finally, we profiled drug induced changes in protein secretion in adult worms to more directly capture treatment associated phenotypes linked to secretory dysregulation. Together, these approaches provide a multidimensional view of antifilarial drug action.

## Results

### Image-based profiling of microfilariae responses to anthelmintics

To establish a baseline for the *in vitro* effects of existing and emerging antifilarial drugs on microfilariae (mf), we quantified the motility and viability of *Brugia* mf exposed to ivermectin (IVM, 50nM-1mM), albendazole sulfoxide (AZS, 5nM-100µM), diethylcarbamazine (DEC, 50nM-1mM), and emodepside (EMO, 500pM-100µM) at 24 and 48 hours post-treatment. High-content imaging data was processed using wrmXpress [35] to generate dose-response curves for parasite motility (optical flow) and to assess viability (green fluorescence) (**Fig 1A**). These data are consistent with previous observations [17,23,24,36] that the *in vitro* motility and viability effects of antifilarials do not fully explain the mechanism of action of drugs used to clear the microfilariae stage (IVM, ABZ, and DEC) (**Fig 1B-D**). While EMO effects on mf motility can be detected at pharmacologically relevant concentrations (IC50 ∼90nM at 24 hrs) [37], IVM elicits effects only at concentrations much higher than experienced in the host (IC50 ∼3µM; plasma C_max_ ∼83nM), and AZS and DEC exhibit no discernible phenotypic effects. Because IVM, DEC, and ABZ are frequently used in combination, we repeated these assays using combined drug treatments to evaluate whether drug interactions or synergies could be detected. The addition of AZS and DEC to IVM treatment or DEC to EMO treatment did not significantly alter the phenotypic responses of mf (**Fig 1E**). Overall, motility results are consistent between *B. pahangi* and *B. malayi* mf across the treatments tested (**S1 Fig** and **S1 Table**). Across all tested drug conditions, paralytic effects are associated with only subtle impacts on tissue viability, reflecting that none of these drugs are directly microfilaricidal [38–40]. However, morphological differences were observed among paralyzed worms treated with IVM and EMO (**Fig 1D**).

**Fig 1.**
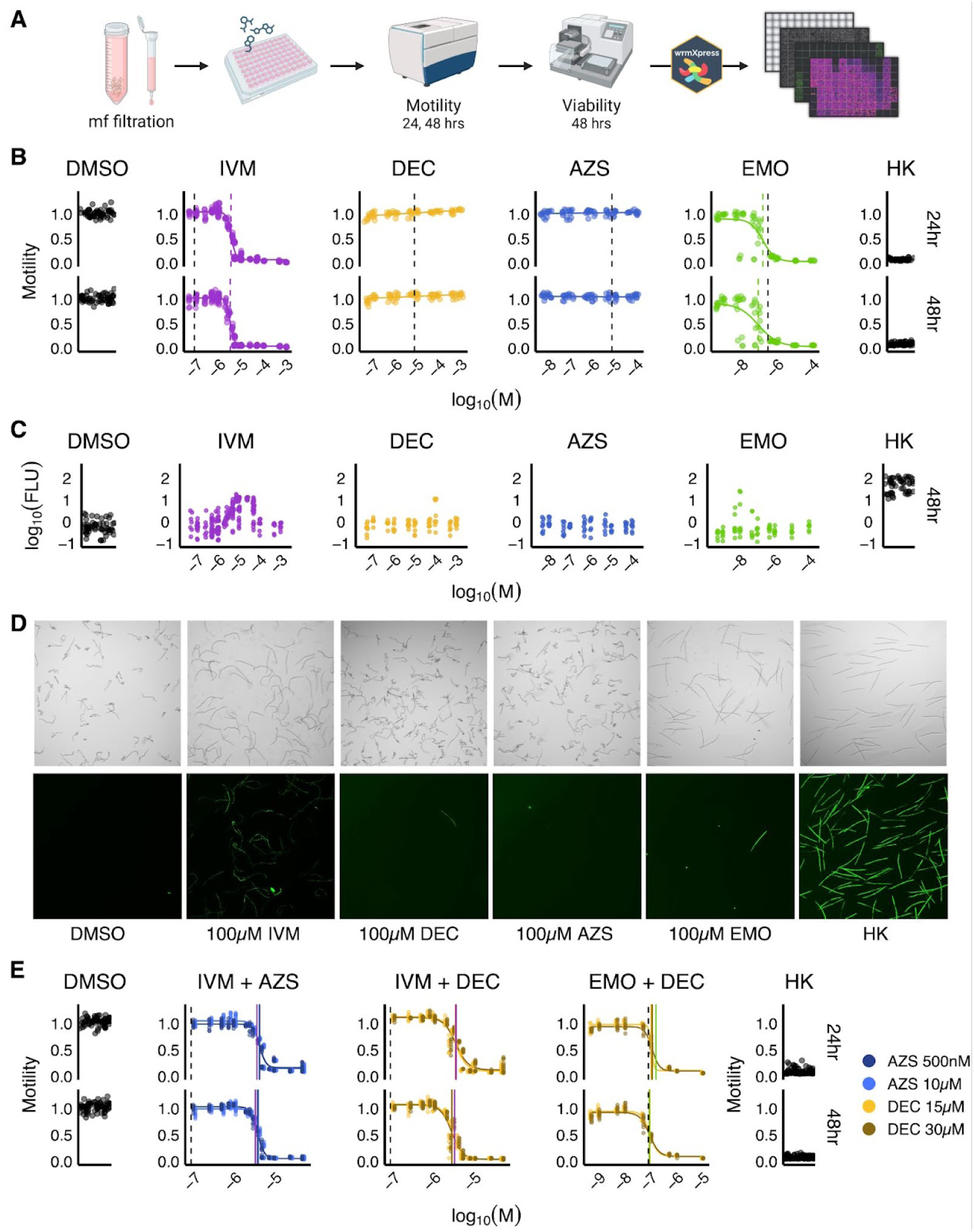
*Brugia* mf motility and viability curves for single and combined drug treatments. **(A)** Schematic depicting the methodology and time points of mf motility and viability data collection. **(B)** Motility dose response curves 24 hours and 48 hours after treatment with ivermectin (IVM), diethylcarbamazine (DEC), albendazole sulfoxide (AZS), and emodepside (EMO), with dashed lines showing experimental IC50 (color) and therapeutic plasma C_max_ (black) values. Controls include mf treated with 1% DMSO and heat killed (HK) mf. **(C)** Viability (CellTox Green) fluorescence readings on a log_10_ scale across treatment concentrations compared to DMSO and HK controls. **(D)** Representative brightfield (top row) and CellTox stained (bottom row) images of control and drug treated mf. **(E)** Motility dose response curves for drug treatment combinations. IVM treatment combined with AZS (500nM or 10µM) or DEC (15µM or 30µM), and EMO treatment combined with 15µM or 30µM DEC. Drug combination IC50s are marked as solid colored lines and IVM plasma C_max_ values as dashed black lines. Individual drug IC50s from (B) are also shown (IVM: purple, EMO: green). Each plot point represents measurements for a plate well containing 1000 mf; each condition was performed across at least four technical replicates (wells) per experiment and each experiment was repeated for at least three biological replicates (parasite cohorts).

### Environmental conditions alter the detection of IVM effects in mf

It has been proposed that host-dependent mechanisms, including changes in parasite secretions, explain the disconnect between *in vitro* and *in vivo* drug responses observed for macrocyclic lactones such as ivermectin and potentially other antifilarial drugs [25,41,42]. However, methods to measure drug-induced secretory dysregulation are low in throughput and require large numbers of parasites. We hypothesized that adjusting *in vitro* culture conditions could sensitize our image based phenotyping approach to detect drug induced motility phenotypes at pharmacologically relevant concentrations. The excretory-secretory (ES) apparatus responsible for mf secretion plays an osmoregulatory role [43–45], and mf undergo shifts in temperature during transmission events that could play a role in priming secretory activity. We therefore tested whether changes in salinity and temperature, cues potentially tied to the physiology and remodeling of the secretory system, modulate observable drug effects.

Previous reports showing that ivermectin (IVM) directly affects mf protein secretion at sub-micromolar concentrations [19–21] informed our selection of IVM concentration (1, 10, 50, and 100 nM) to examine acute effects on mf motility at 1, 2, and 4 hr post-treatment under multiple environmental conditions, including temperature (37°C vs. room temperature [RT], **Fig 2A**) and ionic composition (varying NaCl and KPO₄ concentrations, **Fig 3A**). We first observed an overall decrease in mf motility at RT across all conditions and time points, while mf motility remained relatively stable over time at 37 °C (**Fig 2B**). This shift to RT enabled consistent detection of IVM effects on mf motility at relevant concentrations. Specifically, IVM (10–100 nM) induced a modest but statistically significant decrease in mf motility across time points. These effects were also evident as morphological differences not captured by optical flow-based quantification of motility.

**Fig 2.**
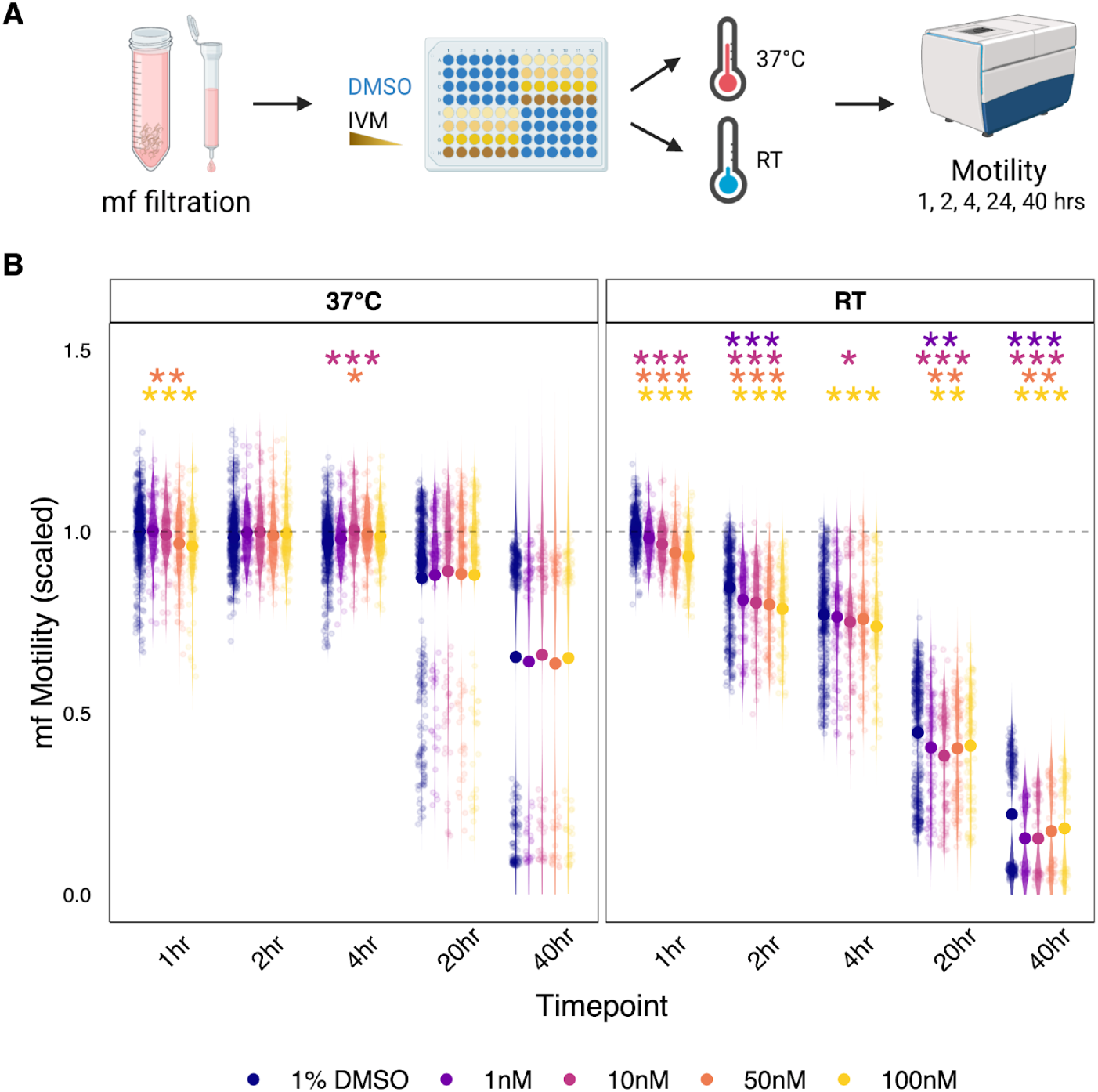
Temperature modulates ivermectin sensitivity of *Brugia* microfilarial motility. **(A)** Schematic showing methodology and timeline for mf temperature shift assay. **(B)** Mean motility, scaled to DMSO 1 hour values of *B. pahangi* mf at 37℃ (left panel) and room temperature (RT, right panel) across time and ivermectin (IVM) or control treatment concentrations (color-coded). P-values represent statistical differences in mf motility between DMSO and drug treatments at matched time points and temperature and were calculated using Anova/Tukey post-test and significance reported as follows, * : p<0.05, ** : p<0.01, *** : p<0.001. Each plot point represents measurements for a plate well containing 1000 mf; each condition was performed across at least six technical replicates (wells) per experiment and each experiment was repeated for at least three biological replicates (parasite cohorts).

**Fig 3.**
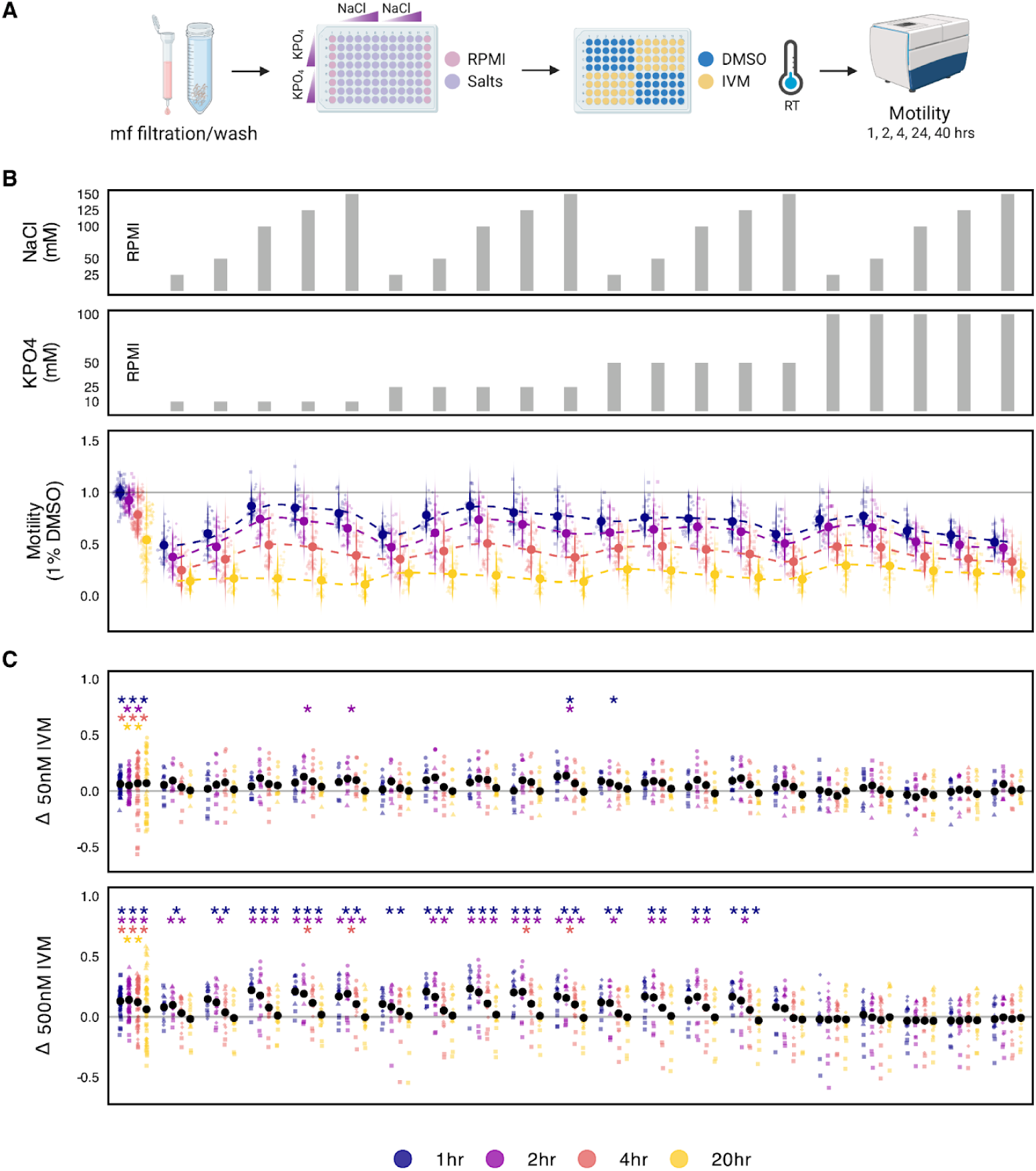
*Brugia* mf motility in the presence of NaCl and KPO_4_ salts. **(A)** Schematic depicting the salt assay methodology and timeline. **(B)** Top two bar graph panels indicate combinations of KPO_4_ concentrations (10mM, 25mM, 50mM, and 100mM) and NaCl concentrations (25mM, 50mM, 100mM, 125mM, and 150mM) across the remaining figure panels at vertically aligned positions. The bottom panel shows DMSO-treated *B. pahangi* mf motility in the presence of different concentrations of NaCl and KPO_4_ across time points. **(C)** The top and bottom panels show optical flow differences between DMSO and ivermectin (IVM) treated *B. pahangi* mf (delta motility) at varying salt combinations in the presence of 50nM (top panel) or 500nM (bottom panel) IVM. P-values representing statistical differences in mf motility between DMSO and IVM treatments were calculated using Anova/Tukey post-test and significance is reported as follows, * : p<0.05, ** : p<0.01, *** : p<0.001. Each plot point represents measurements for a plate well containing 1000 mf; each condition was performed across at least two technical replicates (wells) per experiment and each experiment was repeated for at least three biological replicates (parasite cohorts).

We next altered salinity at room temperature to determine whether ionic stress would further enhance our ability to resolve IVM-evoked phenotypes. Relative to RPMI controls, all tested salt conditions reduced mf motility. Low combined concentrations of NaCl and KPO₄ (<50 mM total) produced a pronounced decrease in mf motility across all time points, whereas higher concentrations of either NaCl or KPO₄ (≥100 mM) paired with lower concentrations of the other salt (<50 mM) had a more modest effect (**Fig. 3B**). Despite these changes, mf displayed substantial tolerance to osmotic variation, remaining motile across a broad range of osmolalities (77–570 mOsm/kg) at all time points (**S2 Fig**). Notably, altering salinity did not improve detection of IVM-induced effects; instead, elevated salinity masked IVM-dependent reductions in motility. Specifically, IVM effects at 50 nM were obscured across all salt conditions, and effects at 500 nM were masked in the presence of high KPO₄ (100 mM) (**Fig. 3C**).

### Age-dependent anthelmintic effects on adult stage parasites

While current antifilarial drugs used in mass drug administration effectively clear circulating microfilariae, they do not cause rapid lethality in adult worms, which can persist for years following treatment. We sought to evaluate the sublethal *in vitro* effects of established antifilarial drugs on adult female (AF) *Brugia* parasites using a common phenotypic platform for quantification of motility and fecundity across time points. This analysis was intended to establish baseline adult responses and benchmark our assay conditions against existing literature, while including emodepside (EMO), an emerging antifilarial with reported adulticidal activity, as a comparator.

We first established an assay for motility and fecundity (protocol A) and then modified culture conditions to improve recovery of excretory–secretory (ES) proteins for downstream quantitative proteomic analysis (protocol B) (**Fig 4A**). This included the removal of serum and phenol red from culture media. Adults are sourced from a jird model of infection with a prepatent period of 3-4 months and are extracted across a wide range of ages reflected by months spent in the Mongolian jird host (**Fig 4B**). Because motility and fecundity results showed no significant differences between protocols for matched treatments, replicates from both protocols were combined to analyze the effects of IVM, DEC, AZS, EMO, and combined IVM-DEC-AZS (IDA) treatment on these phenotypes. IVM and EMO caused sustained motility suppression through the assay endpoint at 48 hrs post-treatment, whereas DEC caused a transient decrease in motility followed by recovery consistent with previous observations [12] (**Fig. 4C**). AZS had no detectable effect on AF motility over the same period. Combined IDA treatment recapitulated both the acute motility decrease associated with DEC and the longer-lasting, dose-dependent inhibition characteristic of IVM alone.

**Fig 4.**
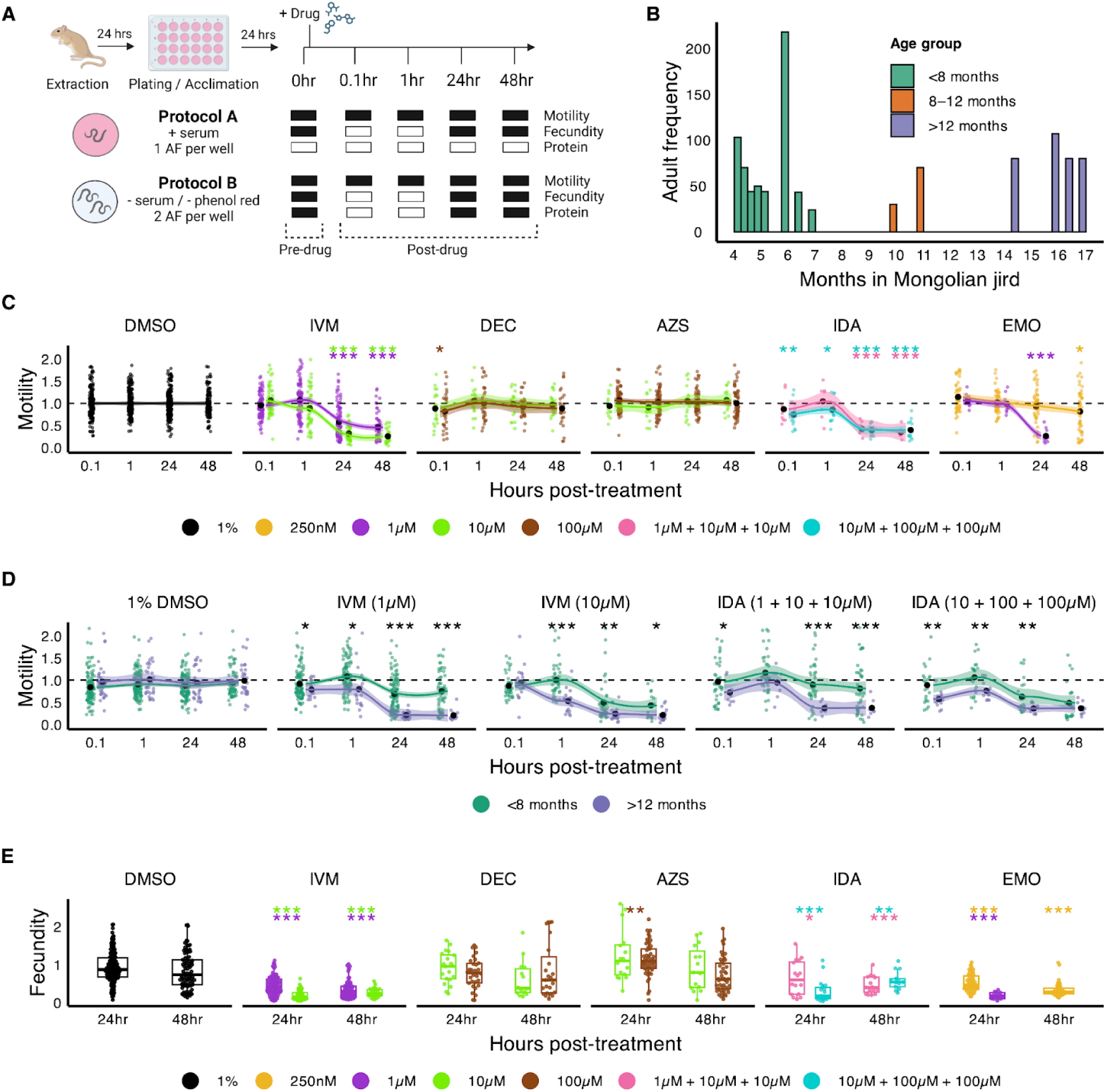
Antifilarial effects on *Brugia* adult motility and fecundity. **(A)** Schematic depicting protocols used to collect motility, fecundity, and protein samples at specific time points for adult female *B. pahangi*. Protocol B was optimized for the collection of excretory-secretory proteins. For all fecundity measurements, media was collected and replaced at 0, 24, and 48 hours. **(B)** Age distribution of adult female worms used in phenotypic assays. **(C)** Motility responses to IVM, DEC, AZS, IDA, and EMO treatments colored by concentration. Motility values were normalized to baseline motility (time = 0) for each individual parasite represented by the black dashed line. Statistical differences are shown for post-treatment time points compared to 1% DMSO controls at the matched time point via t-test to evaluate decreases in motility. **(D)** Motility responses to IVM treatments stratified and colored by the age group adult females occupied. For each drug condition and time point, statistical differences between age groups were calculated via t-test. **(E)** Fecundity responses to drugs as measured by the quantity of progeny released in the presence of drugs after 24 hours and 48 hours, colored by concentration and normalized to 24 hour 1% DMSO controls. Statistical significance was calculated via t-test comparing DMSO and drug treated groups at matched time points. P-values throughout figure are reported as follows, * : p<0.05, ** : p<0.01, *** : p<0.001, **** : p<0.0001. 48 hour motility and fecundity data are not shown for 1 μM EMO as all parasites are paralyzed at this timepoint. Plot points in C-E represent wells of 1-2 adult parasites; each condition was performed for at least 4 technical replicates (wells) per experiment and each experiment was repeated across at least three biological replicates (parasite cohorts).

Although these overall drug response patterns were reproducible across biological replicates, we observed variation in motility sensitivity between parasite batches. These differences correlated with adult worm age, estimated by time spent in the mammalian host. More mature adults (>12 months in host) exhibited increased IVM- and IDA-induced motility inhibition, whereas less mature worms (<8 months in host) showed more variable responses, including partial or complete recovery in some cases (**Fig 4D**). This age-dependent modulation of adult phenotypes, particularly for IVM, represents a previously underappreciated variable.

Adult female fecundity effects were collected by measuring progeny release 24 hours and 48 hours after drug treatment. IVM and EMO inhibited mf release for concentrations tested at both time points, while DEC treatment had no effects on progeny release (**Fig 4E**). Interestingly, AZS treatment induced a small (24%) but statistically significant increase in mf release over the first 24 hours. Combined IDA treatments led to an overall inhibition of fecundity smaller than IVM alone, likely reflecting the opposing effects of IVM and AZS. Fecundity results remained consistent regardless of the time spent in host. Overall, *in vitro* motility and fecundity profiles are complex, with examples of sustained inhibition of motility, recovery, or even enhancement of offspring output. Furthermore, variables like worm maturity impact these results.

### Anthelmintic-induced changes in the adult female secretome

To capture phenotypes more relevant to diagnostics and the host-parasite interaction, we next sought to detect changes in the composition of the adult female (AF) secretome in response to drug. We collected and pooled excretory-secretory proteins (ESPs) at 24 hours and 48 hours post DMSO (1%), IVM (1µM), EMO (250nM) and AZS (100µM) treatments. Media was filtered and proteins (>3 kDa) were concentrated and profiled using NanoLC-MS/MS, resulting in the identification of 88 *B. pahangi* proteins across samples. *B. malayi* orthologs of 55% of these proteins (49/88) were identified in previous adult female proteomic studies [46,47] and dataset comparisons confirm the high abundance of prominent ES proteins, including triose phosphate isomerase (TPI-1), galectin-2 (Lec-2), transthyretin-like family proteins, phosphopyruvate hydratase (Enol-1), cuticular glutathione peroxidase (Bm2151), and macrophage inhibitory factor (MIF-1). Furthermore, we detected 9 proteins previously reported among the 15 most abundant proteins identified in AF extracellular vesicles [20], with four of these proteins (TPI-1, Lec-2, MIF-1, and ACT-5) detected in high abundance in our proteomic dataset. 64% of the *Bpa* proteins were found to have either a classical signal peptide (35%) or an unconventional protein secretion signal (29%), while the remaining proteins were categorized as transmembrane (8%) or intracellular (28%) (**S2 Table**). These results align with previous secretome analyses, which identified classical or unconventional secretion signals in approximately 54%-66% of identified *Brugia* ES proteins from different stages [19,46,47].

The overall distribution of protein intensities across replicates suggests that fewer ES proteins were detected in EMO and IVM treatments compared to DMSO or AZS (**Fig 5A**). Principal component analysis (PCA) shows distinct clustering by treatment; specifically, DMSO and AZS samples grouped together, while EMO and IVM samples form a separate cluster (**Fig 5B**). Normalized protein intensities were used to compare protein abundance across samples and identify differentially expressed proteins (DEPs) for each drug condition compared to DMSO control (**Fig 5C**, **S3 Table**). This identified varying numbers of differentially expressed proteins that met the significance threshold across the three treatment groups. EMO yielded the most extensive list with 29 DEPs (17 up- and 12 downregulated), followed by IVM with 10 DEPs (6 up- and 4 downregulated) and AZS with 5 DEPs (3 up- and 2 downregulated).

**Fig 5.**
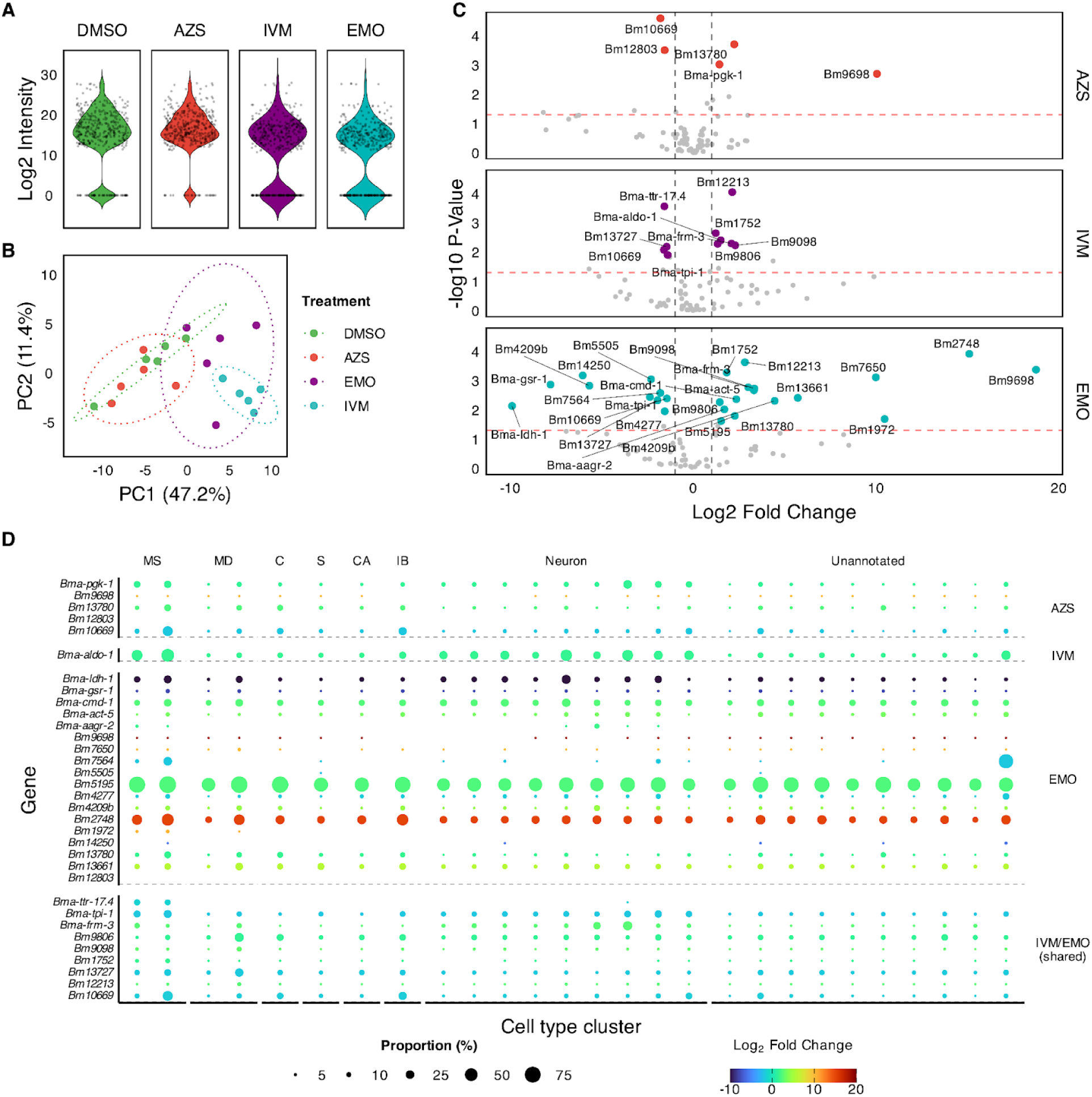
Effects of anthelmintics on the adult female secretory profile. **(A)** Violin plot representing the log_2_ transformed peptide ion intensities distribution of all protein samples analyzed by mass spectrometry. Five biological replicates (reflecting parasite cohorts) were carried out, each representing secretions pooled from 12 adult females (6 wells) for each treatment condition. **(B)** Principal component analysis showing variance among replicates, color-coded by treatment. **(C)** Volcano plots representing the p-value and log_2_ fold change (FC) associated with protein intensities 48 hours post 100µM AZS (red), 1µM IVM (blue), and 250nM EMO (purple) compared to 1% DMSO control. Red and grey dashed lines represent p-value < 0.05 and |log_2_FC| > 1, respectively. Colored points represent differentially expressed proteins of interest (|log_2_FC| > 1, p-value < 0.05, and FDR < 0.05). **(D)** Dotplot representing differentially expressed *B. pahangi* ES proteins post drug treatment mapped to single-cell gene expression patterns of one-to-one orthologs defined using *B. malayi* mf [52]. Cell annotations associated with cell type clusters are shown at the top (MS: muscle, MD: mesoderm, C: coelomocyte, S: secretory, CA: canal associated, IB: inner body), with dot size reflecting the proportion of cells within that cluster expressing the transcript of interest and dot color reflecting log_2_FC of protein abundance compared to DMSO. Differentially expressed proteins unique to each drug condition, as well as those shared between IVM and EMO, are plotted using either treatment-specific log_2_FC (unique) or mean log_2_FC (shared proteins).

A notable degree of overlap was observed between the IVM and EMO datasets with 9 of the 10 proteins identified in the IVM group also dysregulated in the EMO group. This shared signature includes TPI-1, FRM-3, and superoxide dismutase (ortholog of Bm13727). Despite these commonalities, most other DEPs were unique to their respective treatments. The only protein consistently upregulated across all three drug conditions was the cystatin-type cysteine proteinase inhibitor CPI-2 (ortholog of Bm10669). Both SOD and CPI-2 are known to play critical roles in mediating host-parasite interactions [48–51], suggesting that these treatments may trigger common pathways involved in the parasite’s defense against host-induced stress. Finally, we investigated whether ES proteins that are up or downregulated in response to drug are associated with or likely to originate from specific tissues. The expression patterns of proteins are not comprehensively mapped across adult *Brugia*, so we instead mapped proteins of interest to a single-cell RNA-seq atlas produced using *B. malayi* mf [52]. DEPs are generally broadly expressed and likely to originate from a variety of tissues at these sublethal concentrations of drug (**Fig 5D**).

## Discussion

A better understanding of anthelmintic effects on parasites remains critical to improving our ability to treat and diagnose filarial diseases, as well as to the discovery of new drugs. Current *in vitro* assays often fail to capture the pharmacologically relevant effects of drugs used in mass drug administration (MDA). Our initial microfilariae (mf) screening underscored this discrepancy, as ivermectin (IVM), diethylcarbamazine (DEC), and albendazole sulfoxide (AZS) impacted motility at non-pharmacological concentrations, while only emodepside (EMO) recapitulated its efficacy within a therapeutic range. Furthermore, the lack of observable drug synergies suggests that the efficacy of combination therapies like IDA may rely on host-dependent mechanisms rather than direct, additive neuromuscular interference.

This study employed deeper *in vitro* profiling to reveal how environmental variables sensitize parasites to drug action. We demonstrated that temperature significantly modulates mf motility and IVM sensitivity at concentrations better aligned with therapeutic C_max_ values. However, we found that this drug-induced motility difference can be masked by high KPO_4_concentrations. Our working model suggests that elevated extracellular K^+^ may hinder the hyperpolarizing effects of IVM. Because the KPO_4_ application preceded IVM addition in our assays, the resulting depolarization state likely rendered low IVM concentrations insufficient to trigger further physiological shifts, highlighting the importance of ionic context in drug-target engagement.

Since current anthelmintics fail to eliminate adult filarial parasites in humans and animals, we sought to characterize more cryptic drug effects. These experiments revealed that more mature worms are significantly more susceptible to IVM-induced motility inhibition, potentially because older worms are less physiologically fit or because the drugs differentially impact the mf they harbor. We also observed a surprising fecundity effect with albendazole sulfoxide, which appeared to trigger a transient increase in progeny release at concentrations that elicit no motility effects. Conversely, the transient decrease in motility caused by DEC is not coupled to changes in fecundity, further highlighting the independent action of drugs on these phenotypes.

Finally, we used quantitative mass spectrometry to profile shifts in the adult female secretome in response to drug. We found that IVM and EMO treatments cause the dysregulation of several shared and drug-specific secretory proteins, such as SOD and CPI-2, which are expressed across diverse tissue types. Differentially expressed proteins under drug exposure provide new leads that could potentially be leveraged for the development of improved post-drug surveillance tools, provided they are validated in future *in vivo* studies. Ideally, combinations of such markers could be utilized to distinguish between parasite stages, assess reproductive activity, and monitor treatment progress. Developing multi-marker signatures may be necessary to ensure that the presence of surviving, active adult parasites can be accurately detected in elimination zones, offering a much-needed increase in diagnostic resolution for monitoring the success of elimination programs.

## Materials and Methods

### Parasite Shipment and Maintenance

*Brugia malayi* (*Bma*) and *Brugia pahangi* (*Bpa*) adult females (AF) and microfilariae (mf) were provided by the NIH/NIAID Filariasis Research Reagent Resource Center (FR3); morphological voucher specimens are stored at the Harold W. Manter Museum at the University of Nebraska, accession numbers P2021-2023 [53]. Parasites were shipped overnight from the FR3 in RPMI 1640 media supplemented with penicillin/streptomycin (P/S, 0.1 mg/mL). Upon receipt, AF and mf were kept at 37℃ with 5% atmospheric CO_2_ for a 30-45 minute acclimation period before use in assays.

### Drug Sourcing and Stock Preparation

Compounds were sourced as follows: ivermectin (MP Biomedicals, LLC), diethylcarbamazine (MP Biomedicals, LLC), albendazole sulfoxide (Sigma-Aldrich), emodepside (Advanced ChemBlocks, Inc). Stock solutions were aliquoted in DMSO at 100X final concentrations and stored at -20℃ before being thawed for use in experiments. Similarly, stock solutions of 10X NaCl and 5X KPO_4_ were used in mf salinity assays. To make KPO_4_ stocks, 1M K_2_HPO_4_ and 1M KH_2_PO_4_ were mixed to obtain 1M KPO_4_ at ∼pH 7.3 which was used for all subsequent dilutions.

### Microfilariae Motility and Viability Assay

Upon arrival, mf were centrifuged at 800xg for 10 minutes at 20℃ and supernatant was discarded. Pelleted mf were resuspended in 5mL RPMI supplemented with penicillin and streptomycin (RPMI+P/S) and added to a PD10 desalting column (Cytiva) to remove most host cells and parasite embryos following a previously described protocol [52]. Mf were collected and titered to a density of 10 mf/µL (dose response experiments, **Fig 1**) or 14 mf/µL (environmental condition experiments, **Fig 2** and **Fig 3**) to achieve approximately 1000 mf per well.

Drugs and mf were aliquoted to 96-well plates per assay conditions. For dose-response and temperature assays, 1 µL of DMSO or drug stock were added to wells prior to the addition of 100 µL of mf. For dose response assays, positive control aliquots of mf were heat killed at 60℃ for 1 hour before being added to plates. In temperature assays, to compensate for potential motility loss across the plate during data acquisition, treatments were positioned diagonally across plates (**Fig 2A**), and their positions were alternated in experiment replicates. For salt experiments, 10 µL of NaCl and 20 µL of KPO_4_ stocks were added to wells, followed by the addition of 70 µL MilliQ-washed mf. Ivermectin and DMSO were then added to wells and plates were gently shaken. All plates were sealed with breathable strips and incubated at 37℃ with 5% atmospheric CO_2_ (dose response and temperature assays) or at room temperature (RT) in the dark (salt and temperature assays).

At motility timepoints described for each assay, mf were imaged using the ImageXpress Nano (Molecular Devices) following a previously described protocol [54]. The ImageXpress was set to 37℃ and 5% CO_2_ environmental conditions or left at RT according to assay incubation conditions. To assess mf viability at 48 hours post drug treatments in dose-response assays, mf were treated with the CellTox Green kit (Promega) and fluorescence was measured using the ImageXpress as previously described [54]. Motility and viability images were processed using the motility and mf_celltox modules of the wrmXpress [35] software, respectively. Each assay was repeated with at least three separate shipments of mf (biological replicates).

### Adult Female Assay Set Up

The adult assay was implemented following protocols A or B (**Fig 5A**). In protocol A, parasites were shipped in 50 mL conicals tubes containing RPMI+P/S supplemented with 10% FBS. After brief temperature acclimation at 37℃, media was replaced with pre-warmed RPMI+P/S, and adults were allowed to recover at 37℃ in a 5% CO_2_ environment for 30-45 minutes. AF were then transferred to a petri dish, gently untangled, and their fitness was visually assessed based on motility levels and cuticle integrity. Injured or unhealthy worms were discarded and healthy worms of similar fitness were used. Individual AF were transferred to wells of a 24 well-plate containing 750 µL of RPMI+P/S supplemented with 10% FBS. AF were then allowed to acclimate at 37℃ and 5% CO_2_ for 24 hours, prior to drug treatment. Media without FBS supplement was used for the remainder of the assay. A modified version of protocol A (protocol B) was used to collect secreted proteins. In this protocol, FBS-free media was used in all stages, media lacking phenol red was used for all steps post shipment, and two AF were placed into each well. For both protocols, motility and fecundity samples were collected across four (protocol A) or six (protocol B) technical replicates (wells) per condition and repeated at least three times as follows.

### Adult Female Motility and Fecundity Sample Acquisition

#### Motility acquisition

After 24 hours of acclimation, worms were transferred to a new 24-well plate containing pre-warmed media and kept at 37℃ for 10-15 minutes to avoid temperature-dependent motility changes. The first time point (0 hour), was recorded as previously described [55] followed by drug or DMSO addition. Immediate and 1 hour post-treatment video acquisitions were collected (0.1 and 1 hour, respectively). At 24 hours post-treatment, the AF were transferred to a 24-well plate containing media pre-supplemented with drug or solvent and allowed to acclimate for 10-15 minutes at 37℃ with 5% CO_2_ prior to video acquisition (T=24hr). At 48 hours post-treatment, AF motility was recorded and AF were transferred to a petri-dish for disposal.

#### Fecundity collection and progeny quantification

At 0 (acclimation), 24, and 48 hours post treatment, conditioned media from the 24-well plates was collected in individual tubes and centrifuged (800xg) for 10 minutes to pellet progeny. Following centrifugation, 500 µL of supernatant was either discarded or retained for protein analysis. Concentrated progeny in 250 µL of media were preserved at 4℃, and 50 µL aliquots were transferred to 96-well plates and imaged using an ImageXpress Nano as previously described [55]. Motility and fecundity images were processed using a conda optical flow algorithm and Fiji software [56].

### Protein Sample Acquisition

Conditioned media from 6 wells, representing ES products from 12 *B. pahangi* AF were collected at 24 hours and 48 hours post-treatment in individual low-binding protein Eppendorf tubes. Samples were centrifuged to pellet progeny as described above, and 500 µL of supernatant were pooled across technical replicates (6 per treatment), filtered using regenerated cellulose (0.2 µm, Sigma), and stored at -80℃. Samples were thawed and concentrated using a 3kDa centricon centrifugal filter (Millipore-Sigma, Amicon® Ultra Centrifugal Filter), following manufacturer protocol, and washed with phosphate-buffered saline solution (PBS). Concentrated protein samples (∼100 µL) from 24 and 48 hours post-treatment were pooled together for each treatment condition. Five replicates were carried out per drug condition. Resulting samples, representing soluble ES proteins from 12 *B. pahangi* AF over 48 hours post DMSO, IVM, AZS, and EMO treatment were collected and stored at -80℃ prior to mass spectrometry.

### Mass Spectrometry of Protein Samples

#### In-solution enzymatic digestion and Mass Spectrometry analysis

Secreted proteins were concentrated with TCA/Acetone precipitation and subsequently digested with trypsin and LysC proteases as described previously [57,58]. Digested peptides were desalted (Pierce™ C18 SPE 100µl pipette tips) and loaded on Orbitrap Fusion™ Lumos™ Tribrid™ platform using Dionex UltiMate™3000 RSLCnano delivery system (ThermoFisher Scientific) equipped with an EASY-Spray™ electrospray source (held at constant 50°C). Chromatography of peptides prior to mass spectral analysis was accomplished using capillary emitter column (PepMap® C18, 2µM, 100Å, 500 x 0.075mm, Thermo Fisher Scientific) with 46-minute primary gradient from 4 to 20% acetonitrile followed by 16-minute secondary gradient from 20 to 30% acetonitrile which concluded with a rapid 5-minute ramp to 76% acetonitrile and 4-minute flush-out. As peptides eluted from the HPLC-column/electrospray source survey MS scans were acquired in the Orbitrap with a resolution of 120,000 followed by HCD-type MS2 fragmentation into Ion Trap (30% collision energy) under ddMSnScan 1 second cycle time mode with peptides detected in the MS1 scan from 350 to 1600 m/z; redundancy was limited by dynamic exclusion and MIPS filter mode ON.

#### Data analysis

Analysis was performed to establish relative abundances based on identified peptide ion intensities using Proteome Discoverer (ver. 2.5.0.400) Sequest HT search engine against *Brugia pahangi* proteome [59] (NCBI accession GCA_012070555.1, assembly ASM1207055v1) (14,455 total entries) along with a cRAP common lab contaminant database (116 total entries). Static cysteine carbamidomethylation, and variable methionine oxidation plus asparagine and glutamine deamidation, 2 tryptic miss-cleavages and peptide mass tolerances set at 10 ppm with fragment mass at 0.6 Da were selected. Peptide and protein identifications were accepted under strict 1% FDR cut offs with high confidence XCorr thresholds of 1.9 for z=2 and 2.3 for z=3. Strict principles of parsimony were applied for protein grouping. Chromatograms were aligned for feature mapping and ion intensities were used for precursor ion quantification using unique and razor peptides. Normalization was not performed; protein abundance calculations were based on summed peptide abundances and background-based t-testing executed.

### Proteomic Data Processing and Single Cell Data Comparison

Identified *B. pahangi* proteins were searched against the *B. malayi* NCBI proteomic database and *B. malayi* proteins to identify one-to-one orthologs with >80% amino acid identity. Relative abundances were used based on identified peptide ion intensities from all analyzed replicates to conduct proteomic analysis. Raw intensities were log_2_ transformed in R statistical software (v4.2.2) [60] and normalized with cyclic Loess. Data was then analyzed using reproducibility optimized test statistics (ROTS, v1.26.0 [61]). Resulting log_2_FC, p-values and false discovery rate (FDR)-corrected p-values were used to assess differential ESP expression post anthelmintic treatments. Protein sequences were used to determine the presence of signal peptides using the computational tool outcyte [62].

Previously published single-cell transcriptomic data from *B. malayi* microfilariae were sourced from a previous study [52]. The data were filtered to include only untreated cell populations (“utBM”) and genes with a normalized gene expression count greater or equal to two. The R statistical software (v. 4.2.1) [60] and Seurat single-cell software (v. 4.3.0.1) [63], were used to generate a dot plot of transcript expression across annotated and unannotated cell types with overlapped protein expression values from the proteomic data generated here. The percent of cells within a cluster expressing a transcript of interest was calculated using the DotPlot() function in Seurat (dot size).

## Supporting information

S2 Tabe

## Supplementary Figures

**S1 Fig.**
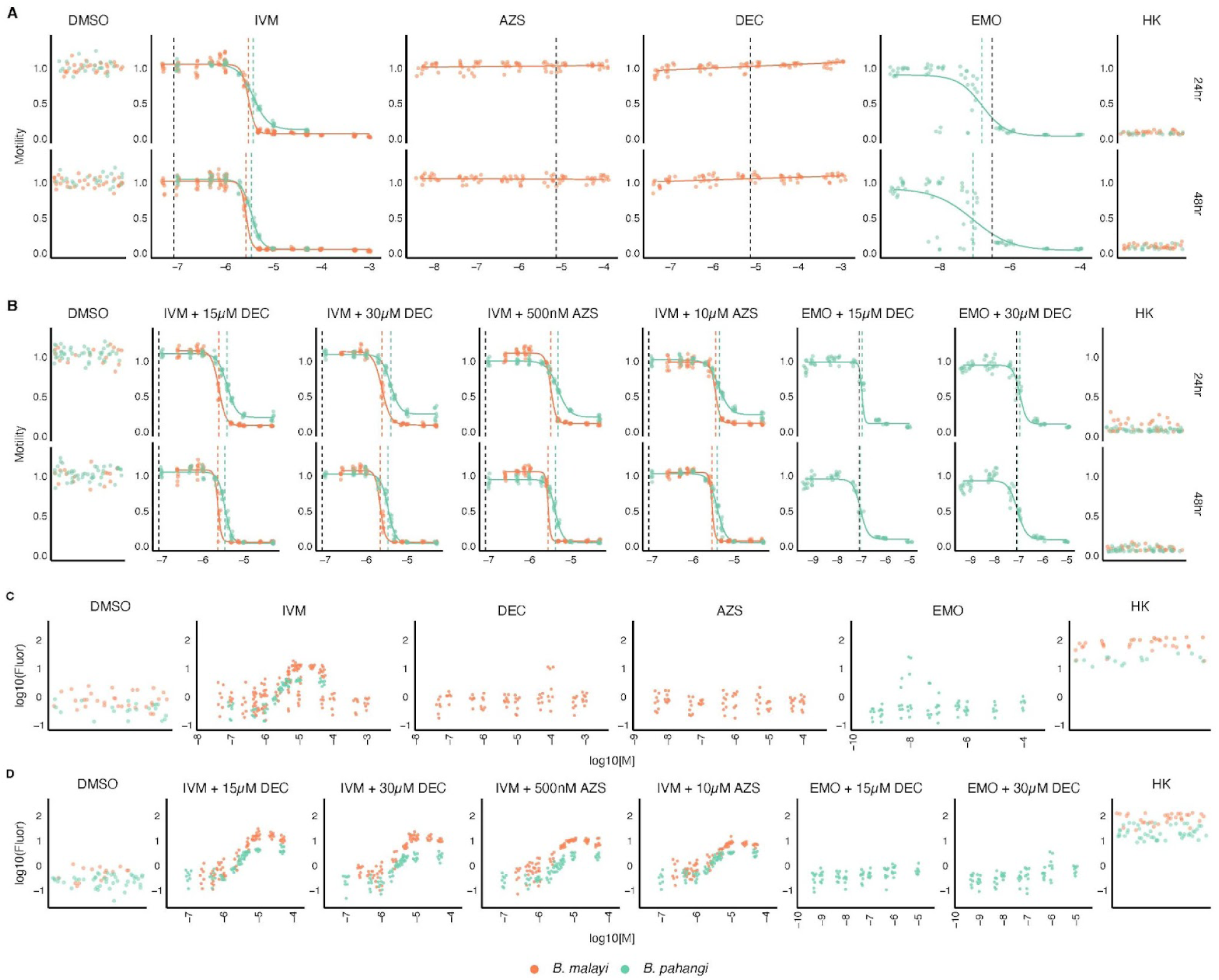
Species-specific dose-responses for mf motility and viability. Panels (A) and (B) display motility curves for single drugs (IVM, AZS, DEC, EMO) and their combinations at 24 and 48 hours, while (C) and (D) show the corresponding viability fluorescence readings. Color represents different *Brugia* species and dashed lines represent experimental IC50 values (colored) and therapeutic C_max_ values (black). DMSO (vehicle) and heat-killed (HK) controls are depicted for each phenotypic assay.

**S2 Fig.**
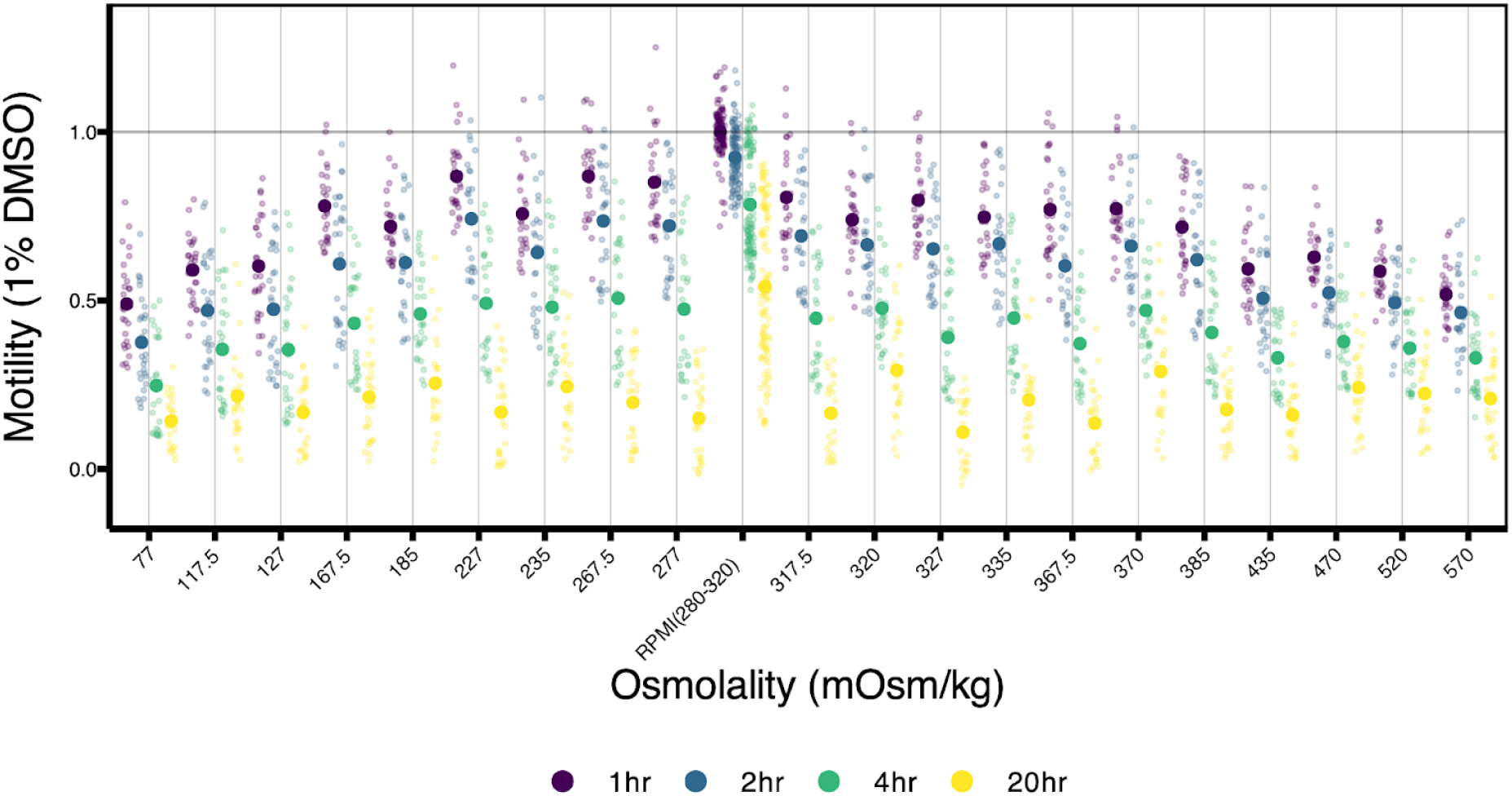
Motility of DMSO-treated *B. pahangi* microfilariae across varying osmolalities. Motility stratified by osmolality calculated for NaCl and KPO_4_ salt combinations. Colors indicate specific time points.

## Supplementary Tables

**<u>S1 Table. IC50 values for microfilariae dose response curves.</u>** Numerical IC50 values for a given treatment and time point calculated for *Brugia* species individually (as shown in S1 Fig) or combined (as shown in Fig 1).

**<u>S2 Table. Adult female ES proteomic data.</u>** Raw peptide ion intensities for all detected *B. pahangi* ES proteins are provided across the five individual replicate drug and vehicle treatments: DMSO, AZS, IVM, and EMO. *B. pahangi* proteins are mapped to one-to-one *B. malayi* orthologs and available gene annotations. Outcyte-based analysis and scores were used to identify proteins with classical signal peptide, unconventional signal peptide (UPS) and to classify the remaining proteins as transmembrane or intracellular proteins.

**<u>S3 Table. Differentially expressed ES proteins.</u>** List of *B. pahangi* proteins and their *B. malayi* orthologs that were identified as differentially expressed in drug conditions compared to DMSO vehicle. Available gene annotations are provided along with log_2_FC, p-values, and FDR derived from ROTS analysis.

## Data Availability Statement

All data and scripts used to process data originating from image-based phenotypic profiling and proteomic datasets are available at IDEA-ms.

## Acknowledgements

This work was supported by National Institutes of Health NIAID grants R01 AI151171 to M.Z. K.T.R. was supported by NIH NIAID grant T32 AI007414. N.J.W. was supported by NIH NIAID Ruth Kirschstein NRSA fellowship F32 AI152347 and NIH NIAID R15 AI183095. We would like to thank Greg Sabat from the UW Madison Biotechnology Center Mass Spectrometry/Proteomics Core Facility for his expertise and valuable suggestions and comments on mass spectrometry data and analysis. Parasite materials were provided by the NIH/NIAID Filariasis Research Reagent Resource Center (FR3). We thank all members of the Zamanian lab for their helpful comments, suggestions, and discussion.

