## Supplementary figures and images for "Deep phenotypic profiling uncovers cryptic effects of antifilarial drugs"

### S1 Fig

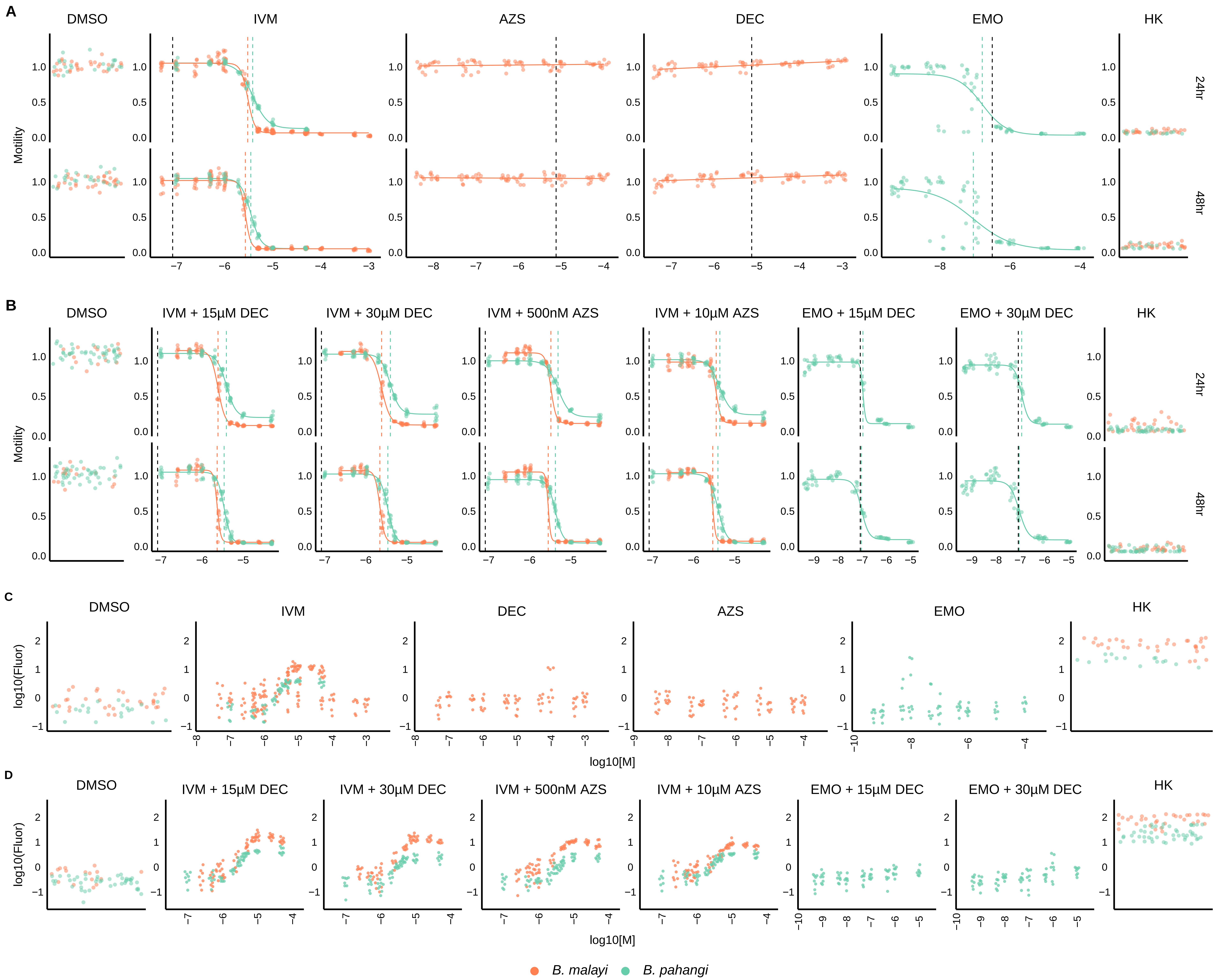

### S2 Fig

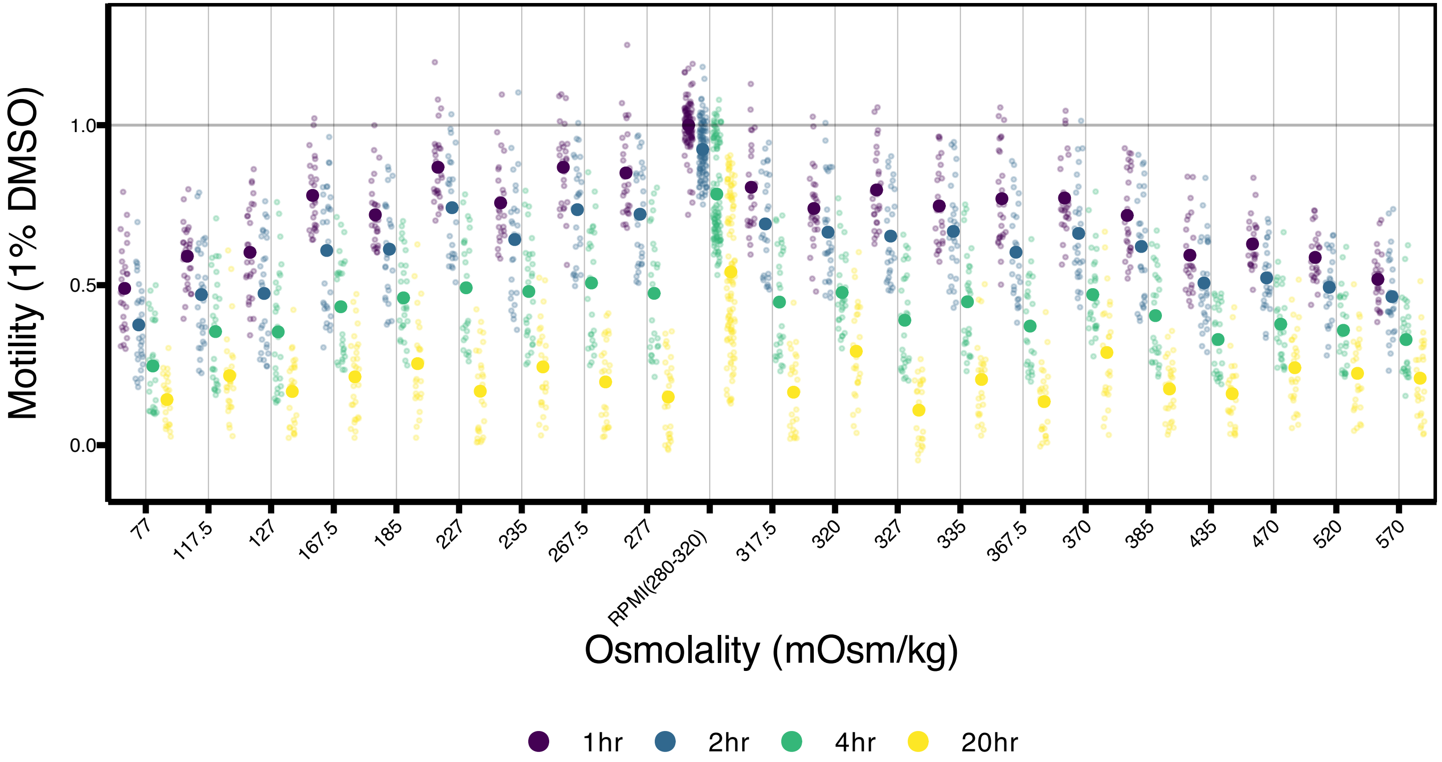
